# Aberrant brain response after auditory deviance in PTSD compared to trauma controls: An EEG study

**DOI:** 10.1101/186460

**Authors:** Katrin A. Bangel, Susanne van Buschbach, Dirk J.A. Smit, Ali Mazaheri, Miranda Olff

## Abstract

Part of the symptomatology of post-traumatic stress disorder (PTSD) are alterations in arousal and reactivity which could be related to a maladaptive increase in the automated sensory change detection system of the brain. In the current EEG study we investigated whether the brain’s response to a simple auditory sensory change was altered in patients with PTSD relative to trauma-exposed matched controls who did not develop the disorder. Thirteen male PTSD patients and trauma-exposed controls matched for age and educational level were presented regular auditory pure tones (1000 Hz, 200 ms duration), with 11% of the tones deviating in both duration (50 ms) and frequency (1200 Hz) while watching a silent movie. Relative to the controls, patients who had developed PTSD showed enhanced mismatch negativity (MMN), increased theta power (5-7 Hz), and stronger suppression of upper alpha activity (13-15 Hz) after deviant vs. standard tones. Behaviourally, the alpha suppression in PTSD correlated with decreased spatial working memory performance suggesting it might reflect enhanced stimulus-feature representations in auditory memory. These results taken together suggest that PTSD patients and trauma-exposed controls can be distinguished by enhanced involuntary attention to changes in sensory patterns.

## 1.0 Introduction

Up to 80% of the general population encounters severe adverse life events. For some, these events can lead to posttraumatic stress disorder (PTSD), a common disorder with a lifetime prevalence of about 7% ^1^^‒^^3^. Diagnostic criteria of PTSD are intrusions, avoidance of distressing trauma-related stimuli, hyperarousal, and negative alterations in cognitions, and mood (see DSM-5^4^) and some patients show attention and memory deficits^5,6^. In particular, the symptom cluster of alterations in arousal and reactivity implies that PTSD patients typically respond to reminders of traumatic events - but also to neutral stimuli - with exaggerated startle response and hypervigilance^4^. High prevalence, difficulties in treatment^7,8^, and often debilitating consequences of the disorder illustrate the great need for investigating the underlying cognitive mechanisms of traumatic stress responses for PTSD. In the current study we investigated whether the brain’s response (measured using EEG) to a simple auditory sensory change was altered in patients with PTSD relative to trauma-exposed matched controls who did not develop the disorder.

The mismatch negativity (MMN) is an event-related potential (ERP) in response to deviant stimuli that differ from a sequence pattern of standard stimuli preceding it^9^. It represents the ability to automatically and without conscious effort compare among series of tones, to detect auditory changes, and to switch attention to potentially important events in the unattended auditory environment^9^^‒^^11^. The MMN provides a neurophysiological index of auditory information processing, perceptual accuracy, learning, and memory; all related to automatic processing and largely not under direct control^12,13^. The MMN plays an important role in clinical research: Weaker MMN can predict psychosis onset^14^^‒^^16^, perceptual and cognitive abnormalities in schizophrenia^17^^‒^^19^, and impaired auditory frequency discrimination in dyslexia^20^. Increases in MMN amplitudes, by contrast, were found in adults with Asperger’s syndrome^21,22^, closed head injury^23^, alcoholism^24,25^, children with major depression^26^, and sleep disorders^27^. Stronger MMN responses for those patient groups may reflect enhanced discrimination of sound pattern^22^ and/or increased involuntary attention switching in response to auditory change^13^.

For PTSD, the MMN remains one of the least studied ERP components and results remain inconsistent (see Javanbakht^28^). Enhanced MMN has been previously found in high school students who have experienced an earthquake^29^ and in women with sexual assault-related PTSD^30^. Those effects may reflect enhanced involuntary attention to auditory deviations in PTSD as a result of chronic hyperarousal and hypervigilance. By contrast, some researchers found reduced MMN in PTSD, which has been interpreted as compensation for chronic hyperarousal^31^ or as difficulty to discriminate relevant from irrelevant stimuli due to overall neural hyperactivity^32,33^. So far, investigations on MMN in PTSD have been limited to specific target groups (high school students) or specific trauma etiologies (earthquake in China) and controls were not matched to the patients. One of our aims is therefore to study same-sex adults diagnosed with PTSD and trauma-exposed controls, matched by age, gender, and education, to investigate group differences in the MMN in response to auditory oddball stimuli.

Event-related potentials such as the MMN reflect change in the EEG over time and are phase locked to the onset of the stimuli. A shortcoming of the MMN as a clinical tool is that it cannot be reliably identified in every individual^34^. As such, another objective of the current study was to examine the induced changes in (i.e. responses in the EEG signal that are time-locked but not phase locked to events) elicited by the onset of deviant stimuli, as a new potential tool for characterizing and predicting PTSD; specifically, the suppression of oscillatory alpha band activity. The depth of stimulus processing in a relevant region is indicative by alpha suppression^35,36^ with the amount of alpha suppression reflecting the resources allocated to processing the stimulus^37^. Stimulus induced alpha modulation in healthy population can be reliably identified within individuals, and covaries with variability in attentional performance ^38,39^.

Investigations into the alpha suppression effect have been applied to research into the neurobiological underpinnings of psychiatric disorders. For example, deviations in alpha suppression have been reported for schizophrenia^40,41^, bipolar disorder, depression^42,43^, and obsessive-compulsive disorder^44,45^. These studies suggest that induced oscillatory alpha activity—even in simple cognitive tasks—may be a valuable biomarker for psychiatry. Here, we will extend current findings and investigate induced oscillatory alpha in PTSD. Research in into oscillatory brain activity in PTSD has been limited to ongoing or resting oscillatory activity. For example, alpha-theta-ratio neurofeedback therapy has shown some degree of effectiveness for Vietnam veterans with combat-related PTSD^46^ and for patients with anxiety disorders, where alpha power changed in proportion to anxiety levels (for review see Moore^47^). Magnetic resonance therapy inducing alpha power increase resulted in decreases in PTSD symptom severity^48^. Finally, alpha asymmetry^49^, alpha peak frequency^50^, and power of right hemisphere frontal alpha correlated with PTSD symptom severity^49,51^ as well as executive task performance^51^.

Few studies have directly investigated alpha power changes in PTSD and results remain inconclusive ^49,^52^,53^. Differences in oscillatory alpha activity during resting state were found among PTSD groups with different trauma etiology. Combat veterans with PTSD were compared to Chernobyl accident survivors with PTSD and showed decreased alpha and increased beta and theta power. However, when compared to controls without the disorder no effects for PTSD were found^54^. Begić *et al*. found increases in the upper alpha, but not in the lower alpha band, in PTSD^55^, while in a later study they found suppression of low alpha over frontal, central and occipital channels^56^. Increased alpha power was found during non-REM sleep for PTSD compared to trauma controls (Olff, personal communication, paper submitted). Besides those resting state EEG studies, a pilot MEG study on nine females with PTSD was conducted and showed reduced alpha power in left Broca area, insula and premotor cortex during tape-recorded auditory trauma imagery compared to neutral imagery^57^. Thus, most research into oscillatory power changes has focused on brain activity during trauma-exposure or at rest. There is strong evidence that hyper-responsivity in PTSD is not restricted to trauma-related stimuli or trauma-related thoughts during rest (see Casada *et al.*^58^vs. Shin *et al*.^59^. PTSD patients show disadvantages in cognitive performance, e.g. tasks that involve attention and memory for neutral information^60,61^. If we detected neural hyperresponsiveness to neutral stimuli that are trauma-unrelated, we could contribute to a better understanding of deviations in automatic attention in PTSD patients. Anomalies in involuntary bottom-up attention in PTSD could explain attention-related problems on everyday tasks.

To investigate this, participants’ EEG was recorded while they were presented with neutral tones according to a simple auditory oddball task while watching a silent movie.

In the current study we investigated whether the brain’s evoked and induced responses to the occurrence of an oddball auditory stimuli were altered in PTSD patients relative to trauma-exposed matched controls. We predicted alterations of the MMN, increased theta band power and stronger suppression of alpha activity after deviant (vs. standard) tones in the PTSD group. We also examined if any of the brain responses correlated with PTSD symptom severity (assessed by the Clinician-Administered PTSD Scale for DSM-4; CAPS^62^) and cognitive task performance (assessed by CANTAB subtests).

## 2.0 Methods

### 2.1 Participants

Thirteen male PTSD patients (mean age = 46.7) were recruited from an outpatient psychiatric clinic. All PTSD patients were diagnosed with chronic PTSD (symptoms duration > 3 months; CAPS score ≥ 45). Thirteen trauma exposed male controls (mean age = 43.5) were matched to the patients based on age and education and met the A criterion (trauma-exposure) for PTSD as defined by the DSM-IV, but no other psychiatric disorders. Controls were recruited from a previous cohort study on prediction and prevalence in PTSD patients from trauma units and through advertising. All participants met the following criteria: no suicidal risk, no neurological impairments, no primary diagnosis of severe depressive disorder, and no other psychiatric disorder (as defined by DSM-IV/ MINI plus^63^), normal hearing, and normal or corrected to normal vision. See also table 1 for participant characteristics. All participants gave written informed consent and received 10 € plus a compensation for their travel expenses. The experiment conformed with World Medical Association Declaration of Helsinki and the study was approved by the local medical ethics committee of the Academic Medical Centre of Amsterdam.

**Table 1.**
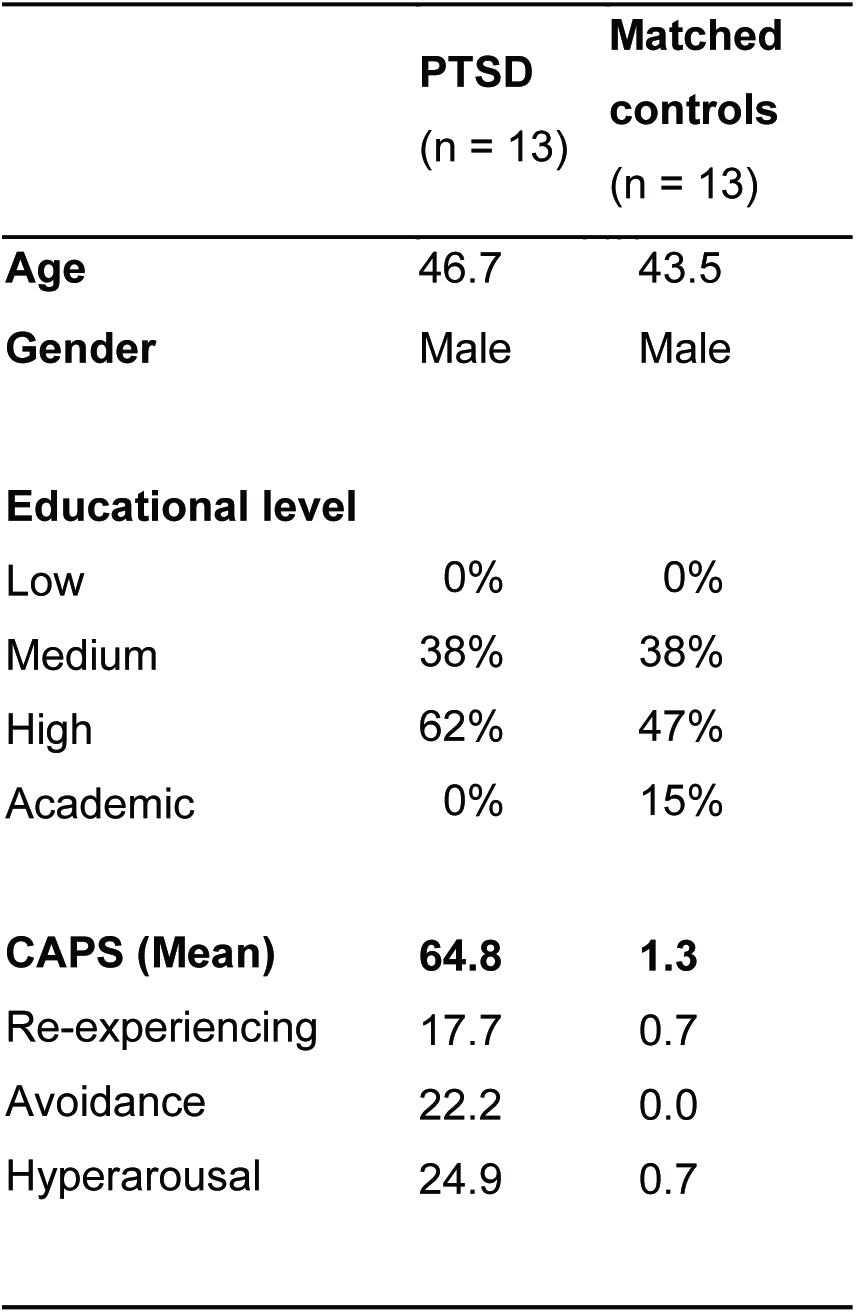
Participants’ characteristics

### 2.2 Auditory paradigm and clinical assessment

The participants were presented with a pseudorandomized sequence of 2000 tone stimuli with a trial duration of 1000 ms (Presentation Software, Neurobehavioral Systems Inc., Albany, CA, USA). The standard tone had a frequency of 1000 Hz and duration of 100 ms. A tone that varied in frequency (1200 Hz, deviant) and duration (50 ms) was added to the sequence and was presented in 11 % of trials but never presented successively. During the presentation of the tones (30 mins), the participants watched a neutral silent movie with subtitles (The life of birds, BBC nature documentary). After the experiment structured interviews were administered after which participants performed additional cognitive tests from the Cambridge Neuropsychological Test Automated Battery (CANTAB). These included the Dutch versions of the M.I.N.I plus to rule out psychiatric disorders^63^, the life events checklist (LEC^64^) to assess potentially traumatic events in a respondent’s lifetime, and the Clinician-administered PTSD scale for PTSD diagnosis and symptom severity assessment (CAPS^65^). Administered CANTAB subtests were the Stop Signal Task (SST) to quantify response inhibition and the Spatial Working Memory (SWM) test.

### 2.3 Electrophysiological Recordings, Pre-processing and Data Analysis

EEG was recorded using a WaveGuard cap system developed by ANT, with 64 shielded Ag/AgCl electrodes (Advanced Neuro Technology B.V.) following the international ‘10/10’ system. To record horizontal and vertical EOG, electrodes were applied to the outer canthi of the eyes and between supraorbital and infraorbital around the right eye. EEG was continuously sampled at 512 Hz with an online average reference and impedances were kept under 100 kΩ. All recordings were done with ASA software (Advanced Neuro Technology B.V.).

Pre-processing of the EEG signal was performed using EEGlab 13.3.2b^66^. The data was resampled to 256 Hz, band-pass filtered between 1 to 30 Hz, and epoched into 1400 ms windows with a 500 ms prestimulus period. For baseline correction an interval of −100 to 0 ms was used. Further analysis of standard tones was restricted to those preceding a deviant tone, since a standard following a deviant tone may be considered deviant. Next, epochs were checked for large artifacts (±75 μV), ignoring high voltage ocular artifacts at frontal pole & anterior frontal locations and visually checked for artifacts. Independent component analysis (ICA) using the logistic infomax ICA algorithm of Bell & Sejnowski^67^ and prior principal component analysis dimension reduction to 30 components. Components containing ocular artifacts were removed and the remaining source-data was back-projected. On average 2.2 and 2.1 components were rejected for controls and patients respectively. Remaining trials that still contained nonbiological artifacts with amplitudes exceeding ±75 μV in any of the channels, were excluded from the analysis and the dataset was visually inspected a last time before entering the statistical analysis. The mean percentage of rejected trials across subjects was 3.2% for controls 4.6% for patients. To investigate the effect of the deviant tone as opposed to the standard tones the datasets were split up into the two experimental conditions again.

#### 2.3.1 Mismatch Negativity Analysis

To analyse MMN, an ERP analysis was performed using the open source FieldTrip toolbox for MatLab (version 20140306)^68^. ERPs were computed for each individual and baseline corrected based on the interval 100 ms prior to auditory stimulus for both trial types separately. Based on the grand-averaged ERPs, a fixed time interval of 100 to 160 ms was chosen to determine the mean peak amplitude of MMN effects. Statistical analysis of the ERP amplitude was conducted on the average of electrodes FC1/ FC2, where the overall mean peak amplitude collapsed across groups appeared to be maximal. We here employed a repeated measures ANOVA with the factors of group (PTSD vs. Control) and tone (standard v. deviant).

#### 2.3.2 Time-Frequency analysis of oscillatory power

To analyse oscillatory power, a time-frequency decomposition of the EEG data was performed to investigate group differences in temporo-spectral dynamics of oscillatory modulations induced by the onset of the deviant stimuli. Using the Fieldtrip software package^68^ function ft_freqanalysis_mtmconvol, time-frequency representations (TFR) of power were obtained per trial (defined as −0.5 s to 0.9 s after stimulus onset) by sliding a single hanning window with an adaptive size of two cycles over the data (ΔT = 2/f). Power was analysed from 2 to 30 Hz in steps of 1 Hz steps for every 10 ms. A difference TFR of deviant vs. standard post-stimulus activity was computed by subtracting the grand average TFR of standard trials from the grand average TFR of deviant trials. Statistical analyses on the difference TFR were conducted separately for theta (5-7 Hz), low alpha (8-12 Hz), and upper alpha (13-15 Hz) frequency bands. The selection of these frequency bands was based on looking at the grand-averaged data (collapsed across groups/conditions) as well prior literature^37,69^. Power from these frequency-bands was subjected to cluster-level permutation tests^70^ (as described below) to statistically assess any difference between the PTSD and control group across all time points from 0 to 500 ms post-stimulus.

The cluster permutation procedure was used to control for multiple comparisons^70^. This procedure uses a standard test (i.e. the two-tailed paired t-tests computed for channel-time pairs) and notes the significance at an arbitrary but fixed level (here, p<0.05). Clusters based on this threshold (i.e. p<0.05) are identified that show significant effects adjacent in space (channels) and time. Spatial clustering used the ‘triangulation’-method, which is based on direction of effect and spatial proximity. For each cluster, the sum of all significant t-values is noted (cluster statistics). Next, the same is done after randomly selecting 1000 permutations of group values (PTSD vs. controls), extracting the maximum summed-t value of the randomly appearing spatial-time clusters (reference statistics). The p-value is based on comparing the observed cluster statistics to the reference statistics. This p-value controls the false alarm rate and was kept below 5% for each significant cluster.

#### 2.3.4 Correlation Analysis

To assess the relation between modulations of the EEG by auditory deviance and cognitive performance we correlated alpha suppression and theta power increase in response to deviant tones averaged over the location and period of interest with cognitive performance on the CANTAB. All values were log transformed to correct for possible data skewness and wide distributions. Then we computed the Pearson’s correlation coefficients of the detected power effects with individual stop signal reaction time and between errors on the spatial working memory task. Due to CANTAB license expiration and missing data of seven participants on the CANTAB tests, the control group was excluded from the correlation analysis.

Using the same procedure, an equivalent analysis was performed to assess the relation between observed EEG effects and PTSD symptom severity by correlating averaged EEG power data with the CAPS scores.

## 3.0 Results

### 3.1 Stronger MMN and late positivity for PTSD in response to auditory deviance

To investigate groups differences in MMN we contrasted ERPs evoked by the onset of the deviant vs. standard tones. Irrespective of group, negative peak amplitudes were more pronounced after deviant tones [main effect Tone: *F*(1,12) = 65.47; *p* < .001, *η*^*₂*^ = 0.85] which confirmed a significant MMN effect for our experimental setup. An interaction effect of Tone x Group revealed a stronger MMN effect for PTSD (M = -1.08μV) compared to their matched controls (M = -0.53μV) [*F*(1,12) = 20.246; *p* < .001, *η*^*₂*^ = 0.63, see Fig. 1].

**Figure 1.**
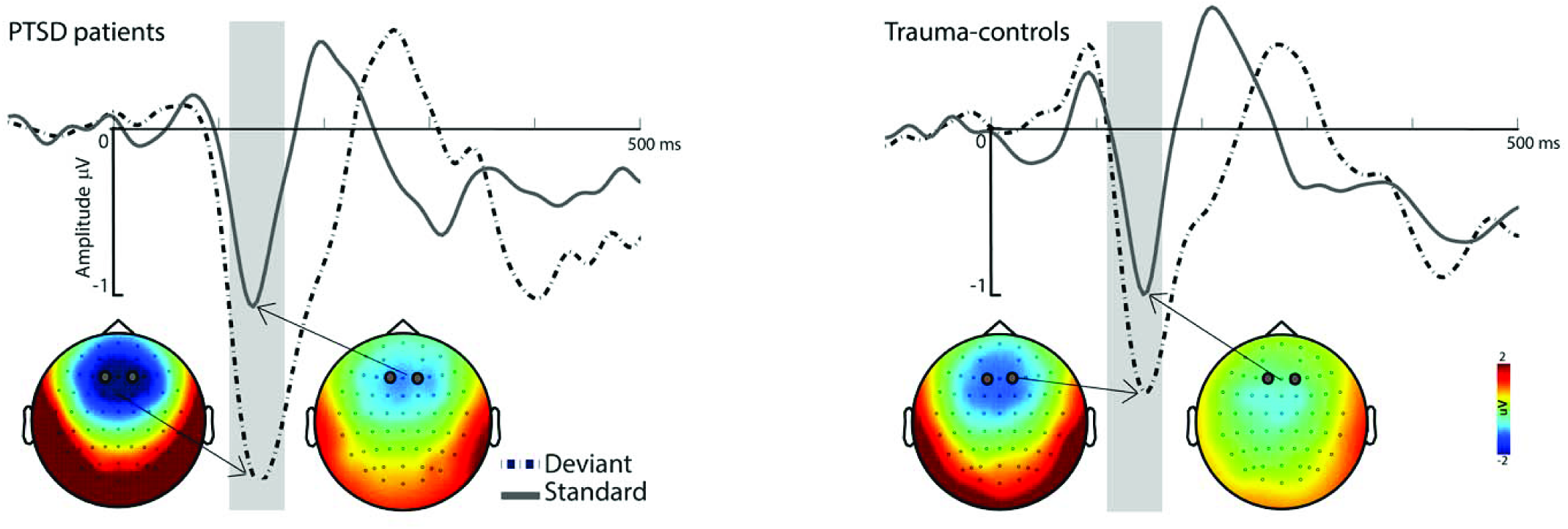
The grand-average ERP waveforms (FC1 / FC2) after deviant and standard tones for PTSD patients (left panel) and trauma-exposed matched controls (right panel). ERP waveforms and topographies show a larger and earlier negative peak (i.e. mismatch negativity) in PTSD patients compared to the controls.

### 3.2 Alpha and theta power modulations

To investigate the differences in oscillatory activity induced by auditory deviance between the PTSD and control groups we subjected the deviant minus standard post-stimulus oscillatory power in the theta and alpha range to a cluster-based permutation test (described above) for all time points between 0 and 500 ms after stimulus onset. Upper alpha power was significantly more suppressed in the PTSD group relative to the controls within the time window of 330 - 490 ms (*p < 0.05*) post stimulus over a cluster comprising right centro-parietal and occipital channels (see Fig. 2). In addition we found that post-stimulus theta power was significantly greater in the PTSD group relative to controls 90 - 250 ms post-stimulus (*p <0.05*, see Fig. 3) over a cluster of frontal electrodes.

**Figure 2.**
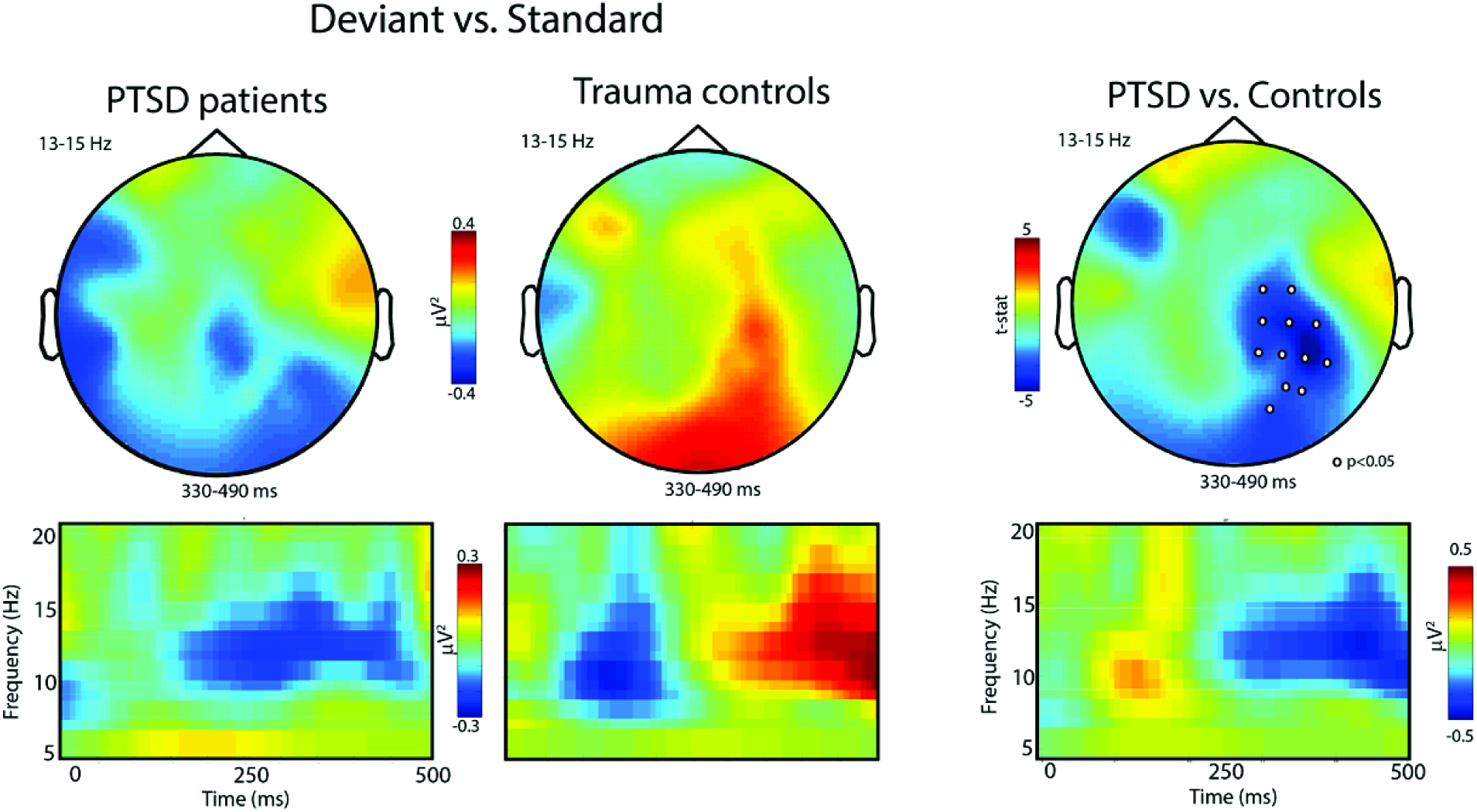
Topographies (upper panel) and time-frequency representations (TFR, lower panel) of upper alpha band power after deviant as opposed to standard tones. Right panel: Greater alpha suppression at 330 – 490 ms after deviant (vs. standard) tones over posterior electrodes in PTSD patients compared to trauma-exposed matched controls suggests hyperresponsiveness to stimulus change, increased auditory processing, and/or stronger stimulus-feature representations in auditory sensory memory in PTSD. Note that topographies were averaged over time window and TFRs over electrodes where a cluster-based permutation test detected a significant group effect in response to auditory stimulus change.

**Figure 3.**
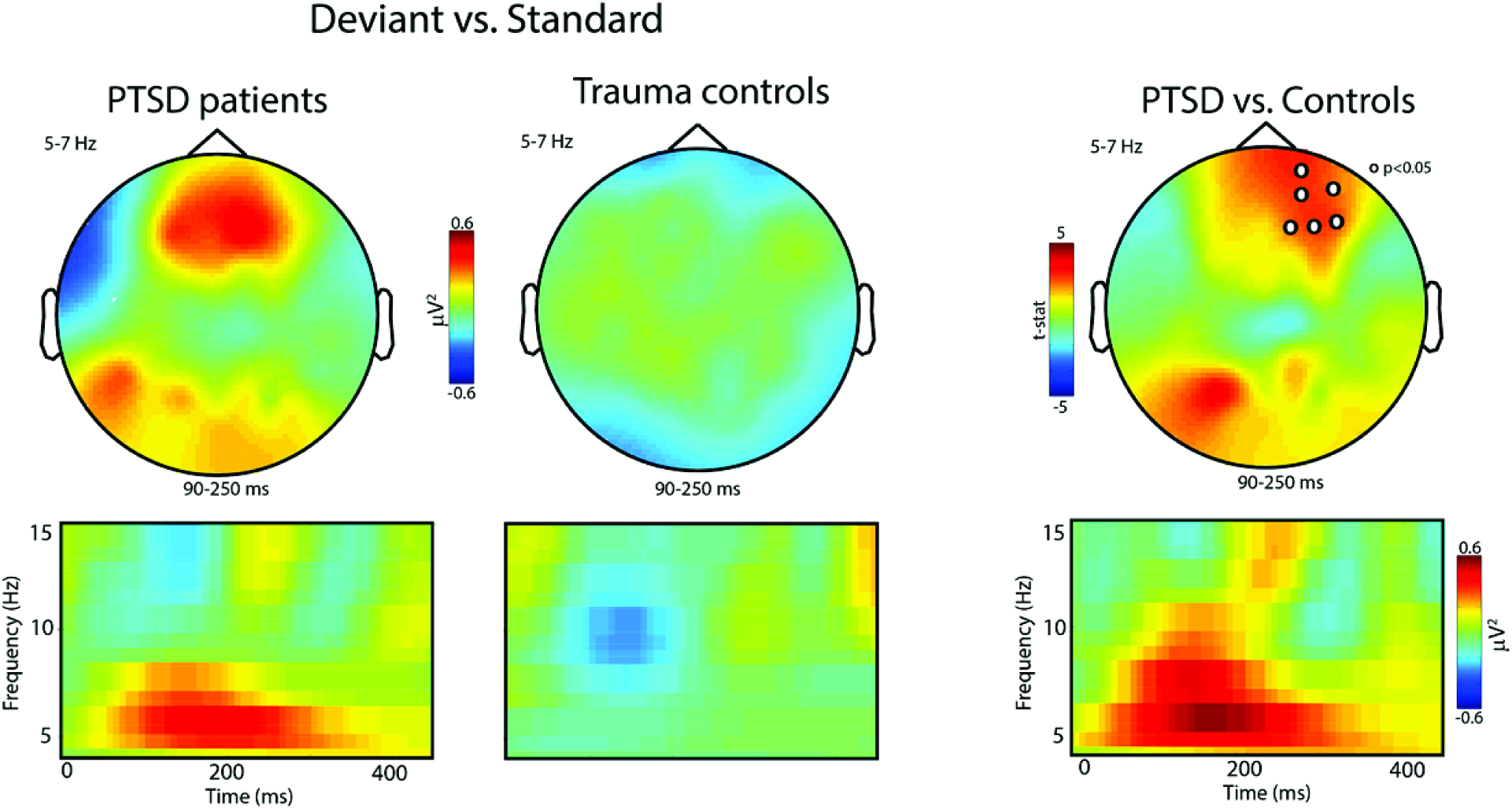
Topographies (upper panel) and time-frequency representations (TFR, lower panel) of theta band power (5 −7 Hz) over frontal channels at 90 – 250 ms after deviant as opposed to standard tones. Right panel: Compared to controls, PTSD patients showed increased synchronization in theta band power over (right) fronto-central channels which indicates enhanced automatic pre-attentive auditory processing and hypersensitive auditory discrimination ability in PTSD in response to deviant (vs. standard) tones.

### 3.3 Alpha suppression effect related to impaired memory performance

Upper alpha power in PTSD in response to the deviant tones correlated negatively with their individual working memory performance (r=- 0.57, N=13, *p* =.035, see Fig. 4) at the time of recording. All other correlations were not significant.

**Figure 4.**
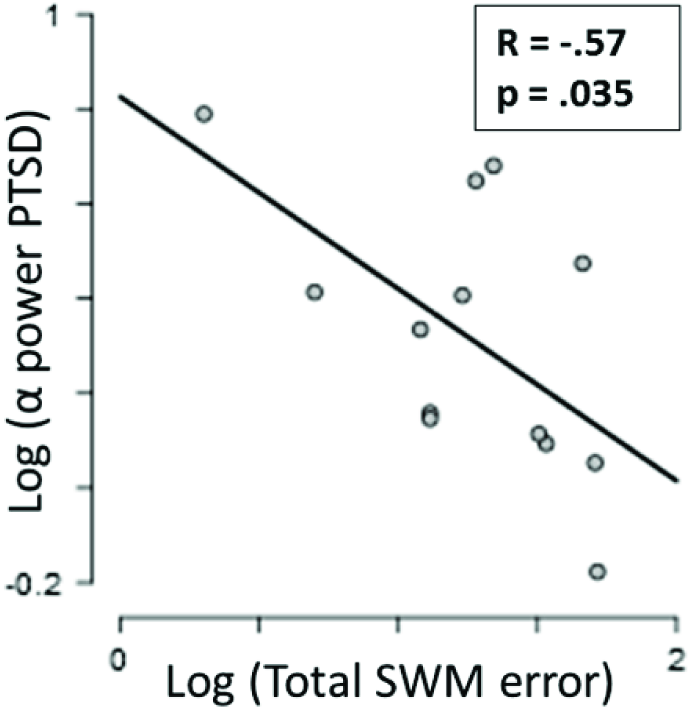
Upper alpha power in PTSD following the deviant tones correlated negatively with patients’ working memory performance. Alpha suppression in response to deviating sounds was related to increasing between errors on the spatial working memory task.

## 4. Discussion

To our knowledge, this is the first study investigating neural responses to deviant vs. standard tones in PTSD patients compared to trauma-exposed gender-, age-and education-matched controls. Participants were presented with an auditory oddball paradigm with 11% of the tones deviating in duration (50 vs. 100 ms) and frequency (1200 vs. 1000 Hz) while watching a silent movie. We found that PTSD patients showed stronger MMN amplitudes relative to controls. In addition, compared to controls, tone deviance induced a suppression of the upper alpha band and increased theta synchronisation in the PTSD group. Alpha power suppression in PTSD was related to impaired individual working memory performance at the time of recording.

Thus far findings of alpha modulations in PTSD have focused on resting state measurements and have remained inconclusive^49,52^‒^56^. Taking these into consideration, while at rest, oscillatory alpha power in PTSD might vary depending on the trauma etiology of the patient group. Patients might either experience the experimental setting as threatening or be more extensively focused on internal processes. Accordingly, in line with our findings, attention to external events would explain results of alpha suppression in PTSD during rest, while inner attention (re-experiencing trauma/ recollection of traumatic memory) would explain findings of increased alpha in PTSD. However, alpha/beta oscillations not only reflect attentional processing, but especially upper alpha band modulations play a role in memory-related processes^71^^‒^^74^. In this study, alpha power suppression induced by deviant stimuli in PTSD was related to impaired working memory performance. Alpha power modulations have previously been observed during attention allocation for memory retrieval and auditory working memory retention which are related to memory performance in healthy participants^75^^‒^^77^. Therefore, given its latency and location, the present auditory-deviance related alpha suppression effect in PTSD might not only reflect attentional mechanisms, but as well account for enhanced auditory stimulus evaluation in PTSD. Patients with PTSD may show enhanced comparison of deviating auditory stimuli to auditory working memory. Since desynchronisation in the upper alpha band has been related to semantic LTM processes^73^ the observed effects even might reflect unconscious automatic retrieval of trauma-related auditory memory in PTSD to compare the salient stimulus with^36,78,79^. This remains to be tested. If true, continuous stimulus comparison to auditory working memory would exhaust and disturb everyday activity that requires memory retrieval which explain the observed correlations of alpha suppression with memory impairment, and the impaired functioning of PTSD patients in daily life.

In sum, increased occipito-parietal alpha suppression and enhanced late positivity after salient tones in PTSD might reflect hypersensitivity to stimulus deviance and abnormalities in the (preconscious) auditory sensory memory. Suppressed alpha power and resulting difficulties in suppression of attentive processing of deviating auditory information could make PTSD patients more prone to automatically detect auditory stimulus change, and to compare those stimuli to auditory short-term memory. This might explain PTSD-specific symptoms of hyperarousal, hypervigilance, memory impairments and distraction from daily tasks.

Our findings of increased frontal theta and increased mismatch negativity in response to stimulus deviance in the PTSD group are in line with previous findings of theta synchronisation in PTSD^55,80^ as well as findings of theta synchronisation in combination with auditory MMN effects in healthy controls. In all three studies those theta effects appeared in about the same time frame and at similar channels locations^81^^‒^^83^. Transient theta as the dominant frequency band during auditory stimulus discrimination has been found to characterize deviant stimulus processing while interfering activations from other frequency networks are minimized^84,85^. MMN amplitudes and frontal theta power have been found to be correlated^86^. Both have been found to be reduced in schizophrenia, which was interpreted as reduced auditory discrimination and shortcomings in prediction and evaluation of stimulus salience^86^^‒^^88^. While auditory discrimination is impaired in schizophrenia^18,89^, PTSD patients might be hypersensitive to auditory stimulus changes. Therefore, the present findings of theta power increases and increased MMN in the PTSD groups most likely reflect hypersensitive early discrimination and change detection processing in the frontal regions. Enhanced automatic pre-attentive auditory processing might explain hypersensitive auditory discrimination ability in PTSD.

The amplitude of the MMN can differ, depending on how prevalent stimulus characteristics between deviant and standard are^90^. The deviant stimuli in the given experiment differed in both, frequency and duration which might explain the early onset of MMN. The average peak onset appears slightly earlier in the patients group (Fig. 1) which could be explained by symptoms of general hypersensitivity/ hyperarousal in PTSD. Mixed results of MMN increase/decrease in PTSD might be explained by the heterogeneity of the disorder. Patients with major symptoms of avoidance (flight-response) may show diminished MMN while patients with stronger symptoms of hyperarousal (like military/police officers) might show increased MMN (see Cornwall *et al.* 2007). This however, remains speculative and more research is needed for clarification.

Limitations of this study would be a rather small sample size that might have limited the power of correlations of EEG effects with PTSD symptom severity (sub-)scores.

Our findings point to a dysregulation in the suppression of task-irrelevant, but salient auditory information in PTSD. While an increased MMN during stimulus perception indicate increased involuntary bottom-up attention switches towards auditory stimulus perception, alpha suppression occurs slightly later and point towards a dysregulation in the suppression of stimulus evaluation at a later stage of stimulus processing. Hyperarousal states can intensify automatic sensory processes that bypass cognitive appraisal and careful evaluation of current perceptions^61^. Here we show that PTSD patients and trauma-exposed controls who do not develop the disorder can be distinguished by their cognitive processing of stimuli that are neutral and unrelated to their trauma. In the search for neuroscientifically-informed treatment interventions targeting specific PTSD symptoms^91^ this study may add to clarifying the neurobiological deviations behind debilitating hyperarousal symptoms interfering with daily functioning in the life of PTSD patients.

## Acknowledgements

A.M. was supported by a VENI grant from The Netherlands Organisation for Scientific Research (NWO).

## Author contributions

A.M. and M.O. designed experiment, S.vB. and K.A.B. performed research, K.A.B. analyzed data and prepared the main manuscript and figures, A.M., D.J.A., M.O. and S.vB. reviewed the manuscript.

## Additional information

The authors declare that they have no competing interests.

